# Oxo-M and 4-PPBP Delivery via Multi-Domain Peptide Hydrogel Toward Tendon Regeneration

**DOI:** 10.1101/2021.08.25.457430

**Authors:** Ga Young Park, Solaiman Tarafder, Samantha Lewis, Soomin Park, Ryunhyung Park, Zain Siddiqui, Vivek Kumar, Chang H. Lee

**Author notes:** **Corresponding author** Chang H. Lee, PhD, Associate Professor, Regenerative Engineering Laboratory, Associate Director, Center for Dental and Craniofacial Research, Columbia University Irving Medical Center, 630 West 168^th^ street, VC12-211B, New York, NY 10032.

## Abstract

We have recently identified novel small molecules, Oxo-M and 4-PPBP, which specifically stimulates endogenous tendon stem/progenitor cells (TSCs) leading to potential regenerative healing of fully-transected tendons. Here we investigated an injectable, multi-domain peptide (MDP) hydrogel providing a controlled delivery of the small molecules for regenerative tendon healing. We investigated the release kinetics of Oxo-M and 4-PPBP from MDP hydrogels and the effect of MDP-released small molecules on tenogenic differentiation of TSCs and *in vivo* tendon healing. *In vitro*, MDP showed a sustained release of Oxo-M and 4-PPBP and a slower degradation compared to fibrin. In addition, tenogenic gene expression was significantly increased in TSC with MDP-released Oxo-M and 4-PPBP as compared to the fibrin-released. *In vivo*, MDP releasing Oxo-M and 4-PPBP significantly improved tendon healing, likely associated with prolonged effects of Oxo-M and 4-PPBP on suppression of M1 macrophages and promotion of M2 macrophages. Comprehensive analyses including histomorphology, digital image processing, and modulus mapping with nanoindentation consistently suggested that Oxo-M and 4-PPBP delivered via MDP further improved tendon healing as compared to fibrin-based delivery. In conclusion, MDP delivered with Oxo-M and 4-PPBP may serve as an efficient regenerative therapeutic for in situ tendon regeneration and healing.

## Introduction

Tendons are dense fibrous tissues with the primary function of transferring mechanical forces from muscle to bone. Injuries to tendons can be caused by laceration, contusion or tensile overload, which account for 50% of all musculoskeletal injuries in the U.S (1-5). For example, rotator cuff injuries affect over 30% of Americans over 60 years of age, leading to over 50,000 surgical repairs annually (6-8). Approximately 11% runners in the U.S. suffer from Achilles tendinopathy (6), and there are 5 million new cases of tennis elbow (lateral epicondylitis) each year (6). This results in a large healthcare burden with treatment for treating tendon injuries exceeding $30 billion per year in the U.S alone (6, 9). Injuries to adult tendons do not spontaneously heal and frequently end up with scar-like tissue - exhibiting high cellularity, disarrayed collagen fibers, and poor mechanical properties (5, 10).

To improve tendon healing, various cell types including tenocytes, dermal fibroblasts and stem/progenitor cells have been applied in tendon tissue engineering *in vitro* or in animal models (9, 11-23). Promising progress has been made in stem cell-based tendon regeneration in vitro and in animal models, despite the lack of clinical availability (18, 19, 24). Recently, we devised a novel *in situ* tissue engineering approach for tendon regeneration by activating endogenous stem/progenitor cells (25). We have identified perivascular-originating TSCs that are capable of guiding regenerative healing of tendons when stimulated by connective tissue growth factor (CTGF) (25). Further investigation into molecular mechanisms of action led us to the discovery of a combination of small molecules, Oxo-M and 4-PPBP sharing intracellular signaling with CTGF, which promotes tendon healing by harnessing endogenous TSCs (26). In addition, our data suggested that Oxo-M and 4-PPBP specifically target CD146^+^ TSCs via muscarinic acetylene receptors (AChRs) and sigma 1 receptor (σ1R) pathways (26). Given no need for cell isolation, culture-expansion and transplantation, *in situ* tendon regeneration by delivery of Oxo-M and 4-PPBP has significant translational potential (27).

Despite a number of advantages (27), small molecule-based regenerative therapies have several limitations. A major outstanding challenge is the fast release of small molecules, likely linked with reduced bioactivity *in vivo* (26). This may serve as a major roadblock in the development of Oxo-M and 4-PPBP as a regenerative therapeutics applicable in large, pre-clinical animal models and humans for which tendon healing likely take a longer than in small animal models (28). Previously, we have investigated efficacy of a controlled delivery of Oxo-M and 4-PPBP via poly(lactic-co-glycolic acids) (PLGA) microspheres (µS) (26). Sustained release of Oxo-M and 4-PPBP from PLGA µS resulted in a significant enhancement in tendon healing (26). However, degradation byproducts of PLGA potentially lower local pH, possibly leading to inflammation and disrupted tissue healing (29, 30). Accordingly, a biocompatible, reliable, injectable and safe vehicle for controlled release of Oxo-M and 4-PPBP is required for facile translation.

In this study, we applied an injectable and self-assembling multi-domain peptide (MDP) hydrogel (31, 32) for controlled delivery of Oxo-M and 4-PPBP. MDP hydrogel is composed of the sequence KKSLSLSLRGSLSLSLKK (termed K2). MDP self-assembles into β-sheets that further form entangled fibrous meshes (31, 32). These highly hydrated meshes generate nanofibrous hydrogels that can be tuned to promote controlled delivery of various bioactive cues (31, 32). Our previous studies confirmed biocompatibility and non-acidic degradation products of MDP (31, 32). Here, we investigated the efficacy of MDP hydrogel with sustained release of Oxo-M and 4-PPBP both *in vitro* and *in vivo* in regard to tenogenic differentiation of TSCs, macrophage polarization and tendon healing.

## Materials and Methods

### Isolation and sorting of CD146^+^ TSC

CD146^+^ TSCs were isolated from patella tendons (PT) of 12 wks old Sprague-Dawley (SD) rats, as per our prior methods (25). Briefly, the harvested PT was cleaned, minced and then digested in 2 mg/ml collagenase at 37°C for 4 hours. After centrifugation of the digest, the pellet was re-suspended in Dulbecco’s Modified Eagle Medium-Low Glucose (DMEM-LG; Sigma, St. Louis, MO) containing 10% fetal bovine serum (FBS; Gibco, Invitrogen, Carlsbad, CA) and 1% penicillin-streptomycin antibiotic (Gibco, Invitrogen, Carlsbad, CA). Then CD146^+^ cells were sorted using a magnetic cell separation kit (EasySep™, StemCells™ Technologies, Cambridge, MA).

### MDP hydrogel for controlled delivery of Oxo-M and 4-PPBP

Multi-domain peptides (MDP) were designed based on previously published sequences: SL: K_2_(SL)_6_K_2_ (32). All peptides, resin and coupling reagents were purchased from CEM (Charlotte, NC). Standard solid phase peptide synthesis was performed on a CEM Microwave peptide synthesizer using Rinkamide resin with 0.37 mM loading, with C-terminal amidation and N-terminal acetylation. Post cleavage from resin, peptides were dialyzed with 500 - 1200 MWCO dialysis tubing (Sigma-Aldrich, St. Louis, MO) against DI water. Peptides were subsequently lyophilized, confirmed for purity using electron-spray ionization mass spectrometry, MicroTOF ESI (Bruker Instruments, Billerica, MA), and reconstituted at 20 mg/ml (20 wt%) in sterile 298 mM sucrose. Gelation of peptide was achieved by addition of volume equivalents of pH 7.4 buffer with 1” PBS or HBSS. Then Oxo-M (10 mM) and 4-PPBP (100 µM) were loaded at 10 – 50 µl in 1 ml of MDP. *In vitro* release profiles were measured by incubating 1 ml of MDP hydrogel encapsulated with Oxo-M or 4-PPBP in PBS or 0.1% BSA at 37 °C with a gentle agitation. The samples were centrifuged at the selected time points, followed by measuring concentrations in the supernatants with a UV-Vis spectroscope (Nanodrop™ 2000, ThermoFisher Scientific, Waltham, MA) at 230 nm and 207 nm wavelengths for Oxo-M and 4-PPBP, respectively.

#### MDP degradation

For *in vitro* degradation test, MDP hydrogel was prepared with and without Oxo-M and 4-PPBP, as labeled with Alexa Fluor^®^ 488 dye. Fibrin gel (50 mg/ml fibrinogen and 50 U/ml thrombin) with and without Oxo-M and 4-PPBP was prepared as a comparison group. The final concentrations of Oxo-M and PPBP in Fibrin gel and MDP were 1 mM and 10 µM, respectively. Then an equal volume (80 µL) of each gel (N = 3) was placed into wells of 24-well plate and kept into PBS for the duration of the study. At pre-determined time points, fluorescent images of the samples were taken using Maestro™ *in vivo* fluorescence imaging system (Cambridge Research & Instrumentation, Inc., Woburn, MA, USA). The images were processed by Image J to calculate percent degradation from the area of the remaining gels.

### In vitro assessment for efficacy of sustain-released Oxo-M and 4-PPBP

Efficacy of sustained release of Oxo-M and 4-PPBP from MDP hydrogel were tested for TSCs differentiation with transwell co-culture. Briefly, MDP encapsulated with Oxo-M and 4-PPBP were applied to Transwell^®^ inserts with 0.4 µm pore membrane, where TSCs (80 – 90% confluence) cultured on the bottom wells. This co-culture model allows transportation of the released small molecules while preventing a direct contact between cells and MDP. At 1 wk culture with tenogenic induction supplements (25), total RNA were harvested and mRNA expressions of tendon related genes including collagen type I and III (COL-I & III), tenascin-C (Tn-C), vimentin (VIM), tenomodulin (TnmD), fibronectin (Fn) and scleraxis (Scx) were measured by quantitative RT-PCR using Taqman™ gene expression assay (Life Technologies; Grand Island, NY) as per our established protocols (25). The quantitative measures for tenogenic differentiation of TSCs by control-delivered Oxo-M and 4-PPBP were compared with release from fibrin gel (50 mg/ml fibrinogen + 50 U/ml thrombin).

### In vivo tendon healing by a controlled delivery of Oxo-M and 4-PPBP

MDP-encapsulated with Oxo-M and 4-PPBP were delivered into fully transected rat patellar tendon (PT) as per prior works (25). Briefly, all animal procedures followed an IACUC approved protocol and 12 wks old Sprague-Dawley (SD) rats (n = 4 per group and time point) were used. Upon anesthesia, a 10-mm longitudinal incision were made just medial to the knee. After exposing PT, a full-thickness transverse incision was made using a no. 11 blade scalpel. MDP hydrogel with or without Oxo-M & 4-PPBP was applied on the transection site. After creating a bone tunnel at the proximal tibia using a 0.5 mm drill, a 2-0 Ethibond suture (Ethicon Inc, Somerville, NJ, USA) was passed through the tibial tunnel and quadriceps in a cerclage technique. The surgical site was then closed using 4.0 absorbable (continuous stitch) for the subcutaneous layer and 4.0 PDS and monocryl (interrupted stitches) for the skin closure. At 2 wks post-op, animals were euthanized and the quality of tendon healing in association with endogenous TSCs were analyzed using H&E, Picrosirius-red (PR) polarized imaging, automated quantitative imaging analysis for collagen fiber orientation. To image whole tissue sections containing any spatial features, slide scanning was performed using Aperio AT2 scanner (Leica Biosystems Inc., Buffalo Grove, IL). From H&E stained tissue sections (n = 10 per group), the quality of tendon healing was quantitatively evaluated using a modified Watkins scoring system (33), covering cellularity, vascularity, cell alignment, amount and size of collagen fibers and wave formation. In addition, immunofluorescence (IF) was performed for macrophage polarization markers, including inducible nitric oxide synthase (iNOS) (PA1-036, Thermo Fisher), and CD163 (NMP2-39099, Novus Biologicals), as co-labeled with DAPI. Anti-inflammatory cytokine IL-10 (AF519-SP, Novus Biologicals) and tissue inhibitor of metalloproteinases-3 (TIMP-3) (ab39184, Abcam) were also evaluated using IF. The labeled tissue sections were imaged using Aperio AT2 scanner with fluorescence.

#### Automated image analysis for collagen alignment

As per our well-established methods (25, 26, 34), we analyzed the collagen fiber orientation in PR stained tissue sections using a digital image processing technique. Briefly, the local directionality and angular deviation (AD) in circularly polarized PR-stained images were calculated by the automated image-processing method. The analysis of each image yielded a distribution of fiber orientations, ranging from -90° to 90°, where 0° was defined as the vertical direction. The degree of collagen fiber alignment was quantified using the AD. The value of the AD was calculated using circular statistics (25, 34) implemented with MATLAB (Mathworks Inc., Natick, MA, USA). For the digital imaging processing, total 15 different image samples were used per group.

#### Modulus mapping with nanoindentation

To assess the maturation and homogeneity of extracellular matrix (ECM) in the healing zone, we performed modulus mapping with nanoindentation on tendon section as per well-established methods (35). Briefly, the nanoindentation was conducted using a PIUMA™ nano-indenter (Optics11, Amsterdam, The Netherlands) with a 1-μm probe. The unfixed and unstained tissue sections were mounted on the embedded high-precision mobile X-Y stage and a maximum force of 10 mN was applied at every 20 μm distance from the original defect site to determine the effective indentation modulus (E_Eff_) across a healed region over selected 400 µm x 400 µm area. The measured E_Eff_ values were displayed in XYZ plane to visualize their homogeneity over unit area. Then the E_Eff_ values from control, fibrin with Oxo-M and 4-PPBP (Fib + OP), and MDP + OP groups were normalized to those of intact region in the corresponding tendon samples.

#### Effect of Oxo-M and 4-PPBP on macrophage polarization

Given the essential roles of macrophages during tendon healing (36), we evaluated effect of Oxo-M and 4-PPBP on macrophage polarization *in vitro*. Briefly, THP-1 human monocytes (ATCC^®^, Manassas, VA) were cultured in complete RPMI media (ThermoFisher Scientific, Waltham, MA), supplemented with 10% heat-inactivated FBS and 1% penicillin/streptomycin (37). For differentiation of THP-1 monocytes into un-activated (M0) macrophages, phorbol 12-myristate 13-acetate (PMA) was applied at 320 nM for 16 hours. For M1 polarization, 100 ng/mL of lipopolysaccharide (LPS) and 100 ng/mL of recombinant human interferon-γ (IFN-γ) were applied for 48 hours. For M2 polarization, 40 ng/mL of recombinant human interleukin-4 (IL-4) and 20 ng/mL of recombinant human IL-13 were applied as per well-established protocols (37). Oxo-M (1 mM) and 4-PPBP (10 µM) were applied along with the M1 and M2 polarization stimuli. After 48 hours, all cells were detached by gentle scraping, followed by RNA isolation for qRT-PCR analysis for M1 and M2 polarization mRNA markers, including tumor necrosis factor alpha (TNF-α), IL-1β, Mannose receptor C-type 1 (MRC1), platelet derived growth factor b (PDGFb).

#### Statistical analysis

For all the quantitative data, following confirmation of normal data distribution, one-way analysis of variance (ANOVA) with post-hoc Tukey HSD tests were used with p value of 0.05. Sample sizes for all quantitative data were determined by power analysis with one-way ANOVA using a level of 0.05, power of 0.8, and effect size of 1.50 chosen to assess matrix synthesis, gene expressions, and structural properties in the regenerated tendon tissues and controls.

## Results

### Sustained release of Oxo-M and 4-PPBP from MDP hydrogel promotes tenogenic differentiation

*In vitro* release kinetics showed that Oxo-M and 4-PPBP are fully released from fibrin within 3 – 4 days (**Fig. 1B**). However, Oxo-M and 4-PPBP showed sustained release from MDP up to 14 - 25 days (**Fig. 1C**). Expressions of tendon-related genes, including COL-I & III, Tn-C, TnmD, Fn and Scx, were significantly increased in TSCs cultured under Trans-well insert loaded with Oxo-M and 4-PPBP in fibrin or MDP hydrogel, in comparison with control with no treatment by 1 wk (**Fig. 1D**) (n = 5 per group; p<0.001). In addition, all the tested tenogenic gene expressions were significantly higher in MDP + OP than Fib + OP (**Fig. 1D**) (n = 5 per group; p<0.001), suggesting positive effect of a prolonged release from MDP hydrogel.

**Figure 1.**
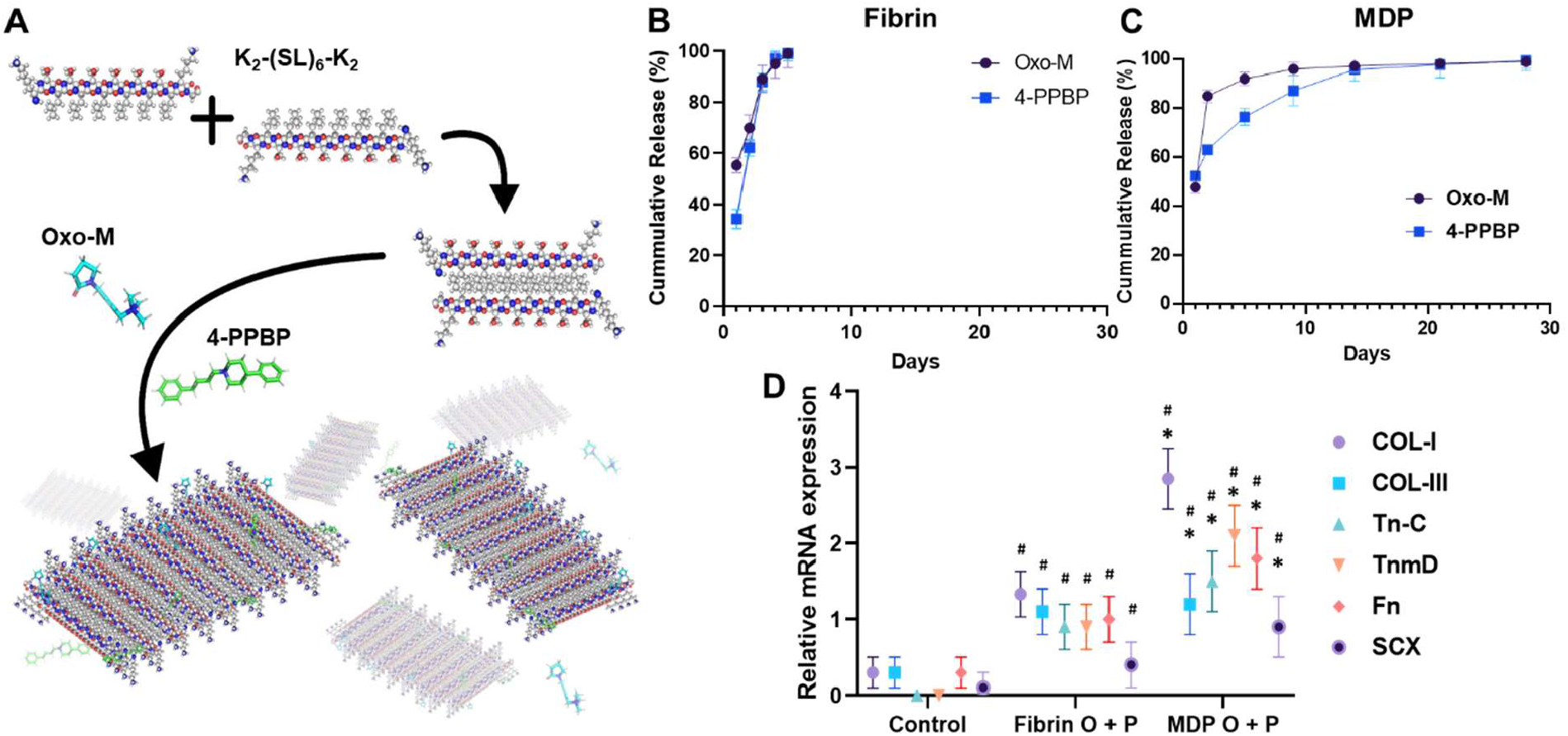
Multi-domain peptide (MDP) hydrogel as controlled delivery vehicle for Oxo-M and 4-PPBP. MDP self-assembles supra-molecularly into nanofibers that encapsulate drugs, while maintaining shear thinning and shear recovery properties (**A**). This allows for facile aspiration and delivery as depots into tissue sites for localized release of small molecule drugs from biodegradable peptide scaffolds. Oxo-M and 4-PPBP loaded in fibrin gel were fully released by 3 – 4 days (**B**), whereas they showed sustained release from MDP hydrogel up to 25 days (**C**). Tenogenic gene expressions were significantly higher in TSCs cultured under Transwell^®^ inserts with fibrin and MDP hydrogel releasing Oxo-M and 4-PPBP (**C**). Oxo-M and 4-PPBP release from MDP hydrogel resulted in significantly higher gene expressions as compared to what released from fibrin (**D**) (n = 5 per group; *: p<0.001 compared to fibrin group; #:p<0.001 compared to control).

### In vitro degradation

Images of fluorescence-labeled hydrogels showed the remaining amount of fibrin and MDP hydrogels over the course of *in vitro* degradation (**Fig. 2A**). Fibrin appeared to fully degrade by 4 days *in vitro* with and without Oxo-M and 4-PPBP. In contrast, MDP hydrogel showed muted degradation by 11 days *in vitro* (**Fig. 2A**). Quantitative fluorescence signal strength measured by Maestro™ imaging system consistently showed that MPD showed ∼48% volumetric reduction by 11 days, which is significantly slower than fibrin gel showing a 100% degradation by 4 days (**Fig. 2B**). In addition, delivering Oxo-M and 4-PPBP in the MDP hydrogel significantly accelerated the *in vitro* degradation (**Fig. 2B**).

**Figure 2.**
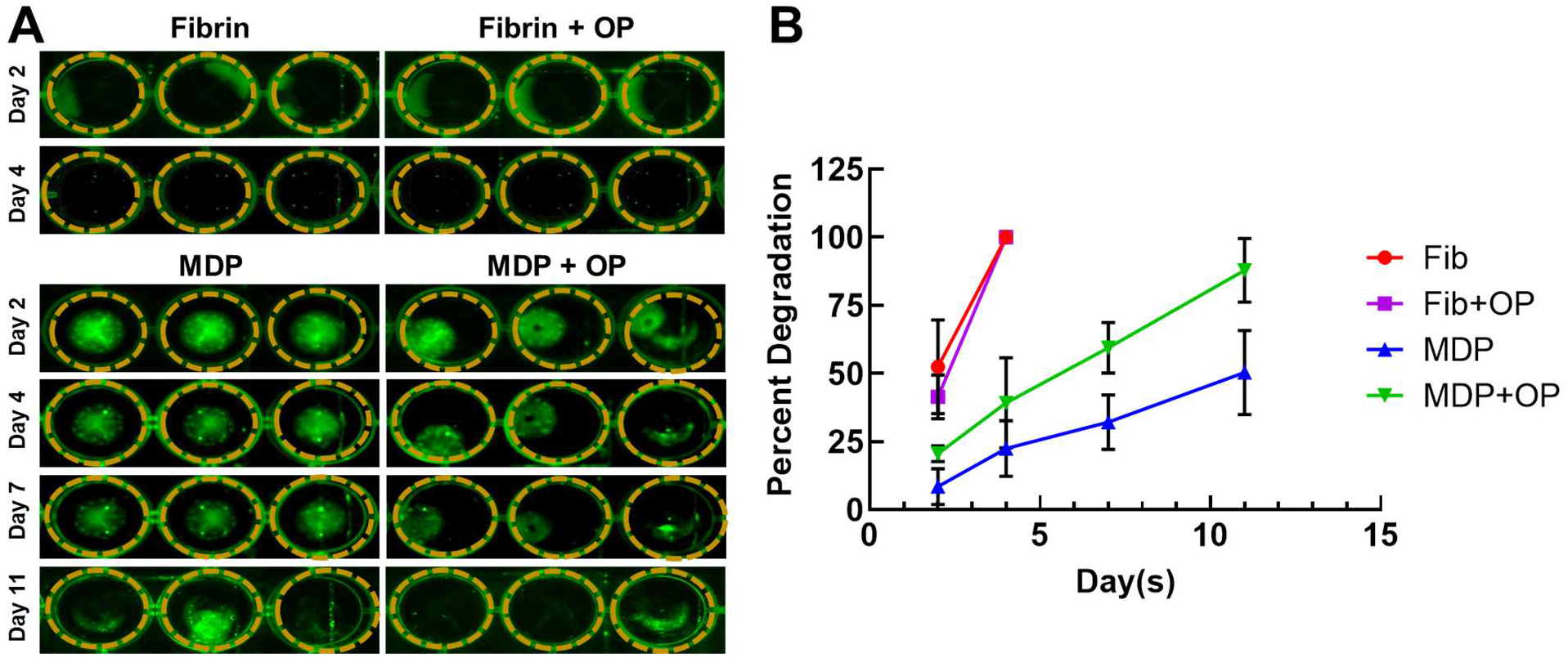
*In vitro* degradation of MDP and fibrin gel with and without Oxo-M and 4-PPBP. Fluorescence-labeled fibrin and MDP with or without Oxo-M and 4-PPBP (OP) were imaged (**A**) and the integral of signal intensities were quantified (**B**) (n = 3 per group).

### MDP delivered with Oxo-M and 4-PPBP enhanced tendon healing in vivo

Fully transected rat PT without treatment ended up with scar-like healing with high cellularity, lacked collagen matrix and disrupted collagen orientation by 2 wks post-op (**Fig. 3A, D, G & J**). In contrast, Oxo-M and 4-PPBP delivery via fibrin and MDP hydrogel significantly enhanced tendon healing with significantly improved structure (**Fig. 3B & E**), dense collagen deposition (**Fig. 3H**), and re-orientation of collagen fibers (**Fig. 3K**) in comparison with control. Few tissue samples in fibrin/Oxo-M and 4-PPBP (Fib + OP) group showed somewhat suboptimal healing (**Fig. 3B**), whereas MDP/Oxo-M and 4-PPBP (MDP + OP) resulted in more consistent healing outcome (**Fig. 3C**). Similarly, the collagen fibers appeared to be denser and better aligned in MDP + OP group as compared to Fib + OP group (**Fig. 3I & L**). In addition, MDP + OP resulted in a significantly higher histological score with smaller variance as compared to Fib + OP with a larger variance (**Fig. 3M**). Consistently, quantitative imaging processing showed that the degree of collagen alignment quantified as AD value was superior with MDP + OP to Fib + OP (**Fig. 3N**) (n = 15 per group; p<0.001).

**Figure 3.**
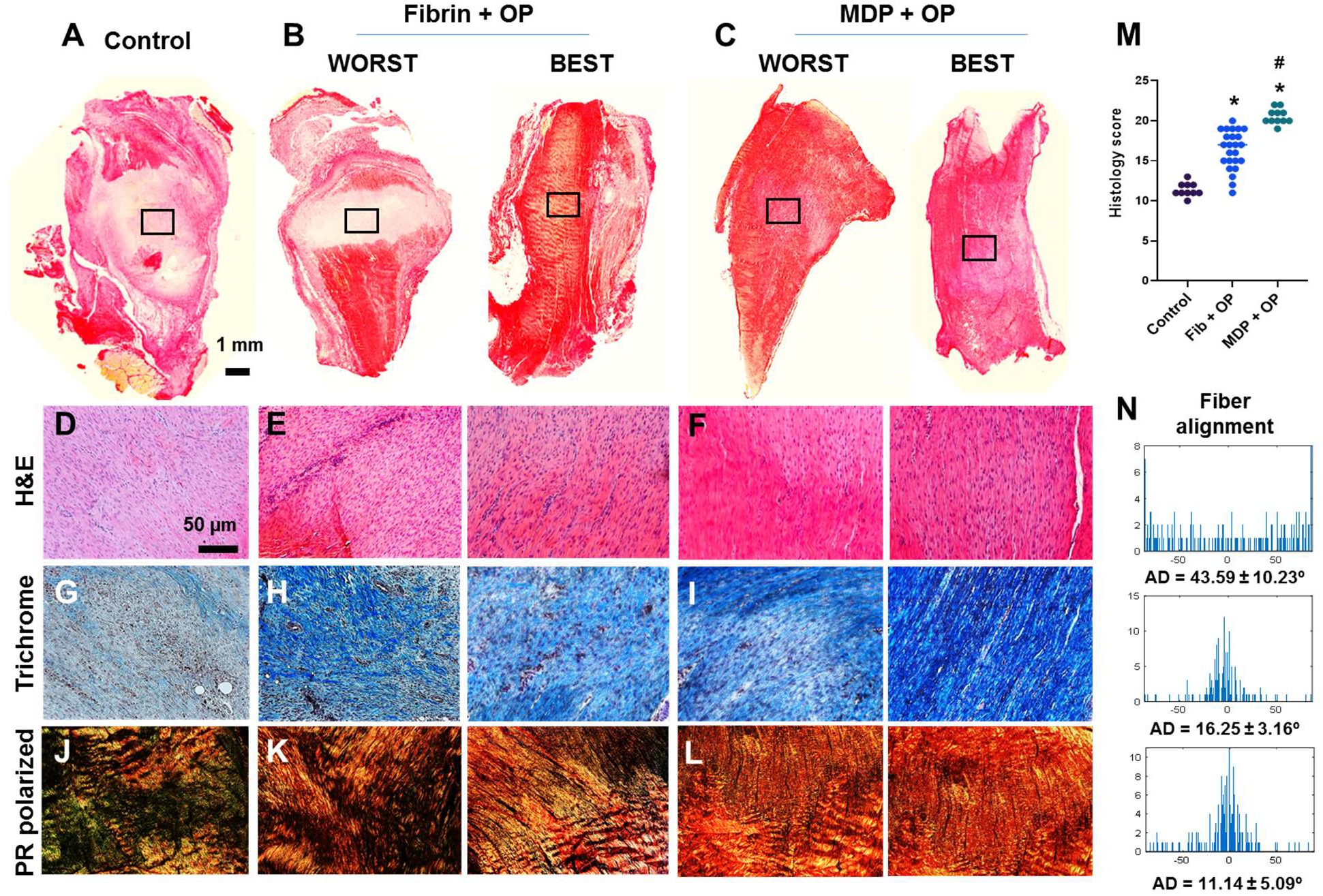
*In vivo* tendon healing by 2 wks. The control ended up with scar-like healing with disrupt matrix and high cellularity (**A, D**), whereas fibrin and MDP delivered with Oxo-M and 4-PPBP showed notable improvement in tendon healing (**B, C, E, F**). Masson’s trichrome showed higher collagen deposition in the healing zone with Oxo-M and 4-PPBP delivery via fibrin and MDP (**G-I**). Polarized PR images showed higher collagen orientation with MDP + OP as compared to fibrin + OP (**J-K**). There were some variances in the healing outcome with fibrin + OP (**B, E, H, K**) in comparison with more consistent outcome with MDP + OP (**C, F, I, L**). Quantitatively, MDP + OP resulted in significantly higher histological scores with a relatively small variance as compared to Fib + OP (**M**) (n = 10 – 25 per sample; *:p<0.001 compared to control; #: p<0.001 compared to Fib + OP). Quantitative angular deviation (AD) value was significantly lower with MDP + OP as compared to fibrin + OP and control (**N**) (p<0.0001; n = 10 – 15 per group). All images are representative best outcome for each group.

### MDP + OP improved magnitude and distribution of indentation moduli

Modulus mapping with nanoindentation displayed the distributions of effective indentation modulus (E_Eff_) over 400 µm x 400 µm areas in the healing regions (**Fig. 4A-C**). Control group showed lower E_Eff_ values with somewhat homogeneous distribution (**Fig. 4A**). Fib + OP showed higher E_Eff_ values with a less homogenous distribution (**Fig. 4B**), and MDP + OP showed a highly homogenous distribution (**Fig. 4C**). Quantitatively, control group showed a lower average E_Eff_ at healing zone than intact tendon, whereas Fib + OP showed average E_Eff_ significantly higher than control (**Fig. 4D**) (n = 100 – 150 per group; p<0.0001). MDP + OP showed E_Eff_ at the similar level with intact tendons (**Fig. 4D**) (n = 100 – 150 per group; p<0.0001). Consistently with E_Eff_ distribution (**Fig. 4A-C**), MDP + OP showed smaller variance in E_Eff_ than Fib + OP (**Fig. 4D**).

**Figure 4.**
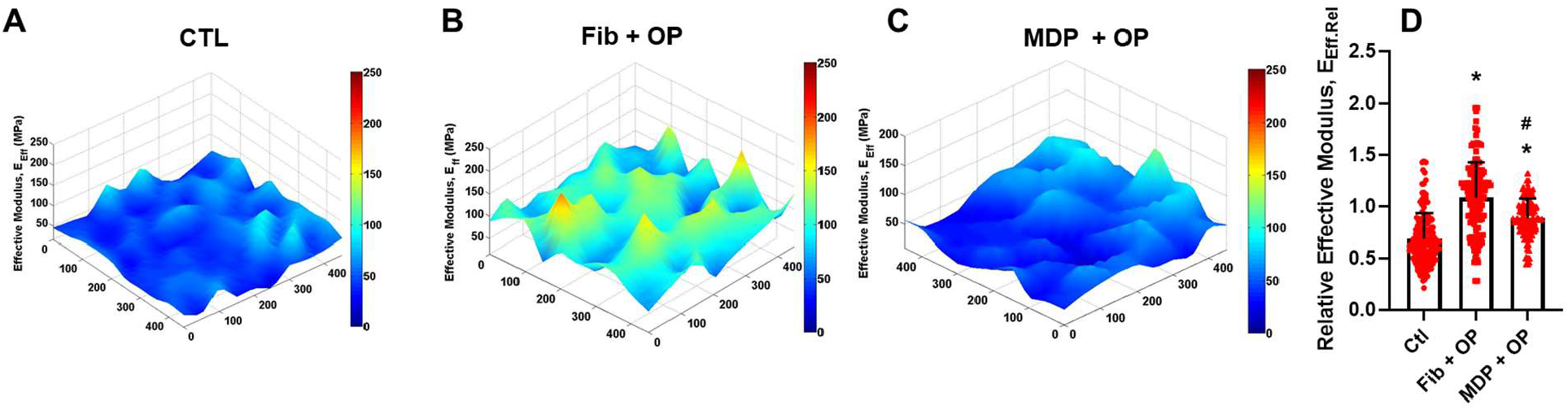
Modulus mapping by nanoindentation of tendon sections (**A-C**), showing more homogenous distribution of indentation moduli with MDP + OP as compared to Fib + OP. Relative effective modulus (E_Eff_.Rel) at healing zone in respect to corresponding intact area (**D**) were significantly higher in Fib + OP and MDP + OP than control. E_Eff_.Rel showed larger variance in Fib + OP than MDP + OP (n = 100 – 150 per group; *:p<0.0001 compared to control; #:p<0.0001 compared to Fib + OP).

#### Effect of Oxo-M and 4-PPBP on macrophage polarization

By 48 hours of M1 polarization of THP-1 derived macrophages induced by LPS and IFN-γ, the treatment with OP significantly reduced mRNA expressions of TNF-α and IL-1β (**Fig. 5A**). In contract, Oxo-M and 4-PPBP significant promoted M2 polarization induced by IL-4 and IL-10, with elevated levels of MRC1 and PDGFb (**Fig. 5A**) (n = 6 per group; *:p<0.001). *In vivo*, OP delivery via fibrin and MDP resulted in a significantly lower number of iNOS^+^ M1-like cells by 2 wks post-op (**Fig. 5B**). The number of CD163^+^ M2-like macrophages was significantly increased with OP delivery (**Fig. 5B**). In addition, MDP + OP showed more M2-like CD163^+^ cells as compared to Fib + OP (**Fig. 5B**).

**Figure 5.**
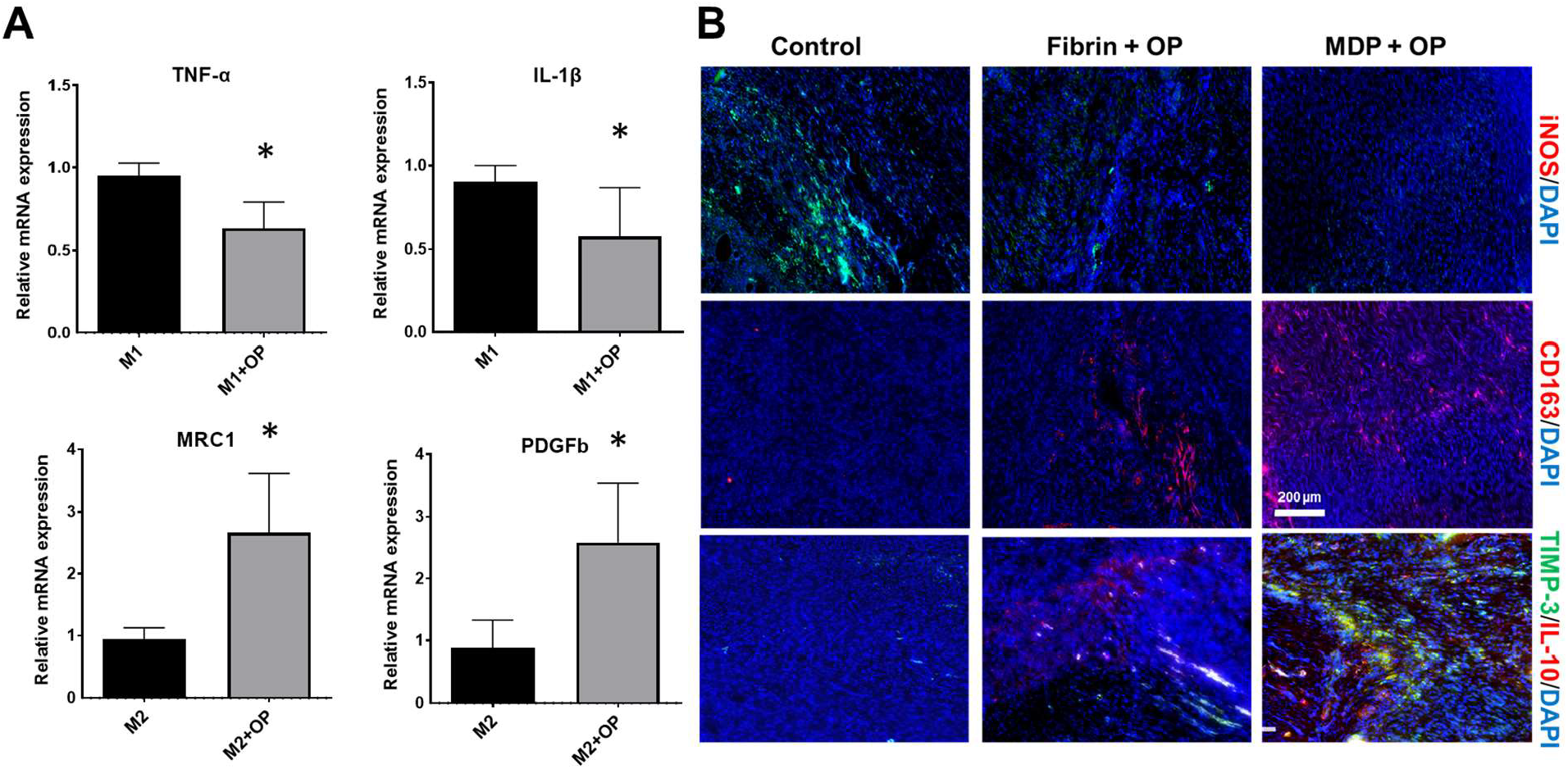
Effect of Oxo-M and 4-PPBP on macrophage polarization (**A**) (*:p<0.001 compared control). Immunofluorescence of macrophage and anti-inflammatory markers (**B**). The number of iNOS+ M1-like macrophages were lower with OP delivery (**B**). MDP + OP showed an increased number of CD163+ M2-like cells as compared to fibrin + OP (**B**). TIMP-3 and IL-10 showed robust expression in MDP + OP group in comparison with Fib + OP (**B**).

## Discussion

Our findings suggest an effective and reliable approach to enable a controlled delivery of small molecules that improve regenerative tendon healing by harnessing endogenous stem/progenitor cells. The unique chemical characteristics of MDP hydrogels, self-assembling into β-sheets, enable entrapment of small molecular weight drugs such as Oxo-M and 4-PPBP, consequently providing sustained release over time. Given that MDP self-assembles through noncovalent interactions of alternating hydrophobic leucine residues and hydrogen bonding of hydrophilic serines (32), both hydrophilic Oxo-M and hydrophobic 4-PPBP were able to be loaded into MDP β-sheet and then showing sustained release without notable difference in the release kinetics between Oxo-M and 4-PPBP (**Fig. 1B & C**). In contrast to the previously used PLGA µS, MDP’s degradation byproducts do not change local pH with a good biocompatibility established in a number of prior studies (32). In addition, its unique near instantaneous self-assembly in aqueous solution allows drug solublization and facile injection of MDP hydrogels in desired sites via a syringe needle, followed by near-instant *in situ* gelation (32). These characteristics further advocate the potential of MDP hydrogels as an efficient controlled delivery vehicle.

A prolonged release of Oxo-M and 4-PPBP from MDP appeared not only to enhance tenogenic differentiation of TSCs but also to modulate polarization of macrophages. Our *in vitro* data suggest that Oxo-M and 4-PPBP may interfere with M1 polarization while promoting M2 polarization. Collective experimental evidences in several previous studies support the temporal roles of inflammatory of M1 macrophages and anti-inflammatory M2 macrophages in the early and late phases of tendon healing, respectively (35, 38-40). Excessive or prolonged M1 macrophages are closely involved with inflammation and scarring, whereas M2 macrophages play essential roles in matrix synthesis and remodeling (35, 38-40). Thus, prolonged activities of Oxo-M and 4-PPBP via controlled delivery with MDP may have promoted tendon healing by attenuating M1-mediated inflammation and M2-mediated anti-inflammatory cytokines and matrix remodeling. Consistently, we have observed the elevated levels of TIMP-3 and IL-10 with MDP + OP as compared to Fib + OP by 2 wks post-op.

The modulus mapping on sectioned tendon tissues by nanoindentation revealed interesting features on healed ECM (**Fig. 4**). Scar-like matrix formed in control group showed a relatively homogenous distribution of indentation modulus with high moduli at isolated area (**Fig. 4A**). Tendon tissue healed with OP delivered via fibrin gel increased the average indentation modulus but showed substantial inhomogeneity over the testing area (**Fig. 4B**). Notably, tendon delivered with MDP + OP resulted in increased indentation moduli with highly homogenous distribution (**Fig. 4C**). These observations may suggest that relatively inhomogeneous matrix in Fib + OP is likely due to immaturity of healed tendon matrix, and more mature tissue matrix in MDP + OP group was formed by a prolonged release of OP leading to sustained activation of M2 macrophages modulating inflammation and matrix remodeling.

Despite the promising outcomes, our study has several limitations that includes the unknown *in vivo* degradation rate. Most biodegradable materials exhibit *in vivo* degradation rates markedly variant from well-controlled *in vitro* studies (41, 42), likely associated with dynamic changes in the biochemical environment *in vivo* affected by inflammation, cell metabolism, and co-morbidities (41, 42). Thus, the actual degradation of MDP hydrogel and consequent release of Oxo-M and 4-PPBP may differ from the *in vitro* data. Nonetheless, such *in vivo* factors are speculated to affect degradation of both fibrin and MDP, consequently validating our comparative *in vivo* study between the two different delivery vehicles. Various state-of-art imaging modalities are being developed to track *in vivo* degradation and release via non-invasive measurements (43-45), which will likely serve as an efficient tool to further optimize delivery vehicles in follow-up studies.

In conclusion, MDP may represent a highly efficient, injectable hydrogel system allowing controlled delivery of Oxo-M and 4-PPBP with specific function to stimulate endogenous stem/progenitor cells and modulate macrophages toward tendon regeneration. Given no need for cell translation, our approach with MDP releasing Oxo-M and 4-PPBP has significant clinical impact as a highly translational approach to induce regenerative healing of tendons.

## Acknowledgements

We thank Aryan Mahajan for creation of the schematics in Figure 1A.

## Author contributions

G.P. was responsible for the primary technical undertaking and conducted the experiments. S.L. assisted nanoindentation modulus mapping. S.P. and R.P. assisted histomorphological analysis. S.T. performed *in vivo* animal surgeries, digital imaging processing and nanoindentation experiment. Z.S and V.K. are responsible for synthesis and purification of peptide hydrogel. C.H.L. is responsible for the study design, data analysis and interpretation, and manuscript preparation. All the authors edited the manuscript.

## Data and materials availability

All the data presented in this study will be available upon request.

